# A method to keep the excitation light sheet in focus in selective plane illumination microscopy

**DOI:** 10.1101/222497

**Authors:** Liang Gao

## Abstract

Keeping the excitation light sheet in focus is critical in selective plane illumination microscopy (SPIM) to ensure its 3D imaging ability. Unfortunately, an effective method that can be used in SPIM on general biological specimens to find the axial position of the excitation light sheet and keep it in focus is barely available. Here, we present a method to solve the problem. We investigate its mechanism and demonstrate its performance on a lattice light sheet microscope.

## Introduction

Selective plane illumination microscopy (SPIM) has become one of the most important 3D live fluorescence imaging techniques after its reintroduction a decade ago (Huisken et al., 2004). In SPIM, an excitation light sheet is used to illuminate the sample from a direction orthogonal to the detection optical axis, so that only the sample in focus is illuminated during imaging. The confined illumination by the excitation light sheet not only gives SPIM the advanced 3D live imaging ability, but also reduces the photodamage and photobleaching significantly at the same time (Keller et al., 2008; Wu et al., 2011; Planchon et al., 2011; Chen et al., 2014).

In theory, the 3D imaging ability of SPIM is only determined by the intensity profile of the excitation light sheet used to illuminate the sample with a given detection numerical aperture (NA). Nevertheless, it also relies on how well the excitation light sheet is placed in focus in practice, as both the 3D spatial resolution and the optical sectioning ability of SPIM decay quickly when the illumination light sheet deviates from the detection focal plane (Gao et al., 2014, 2015). The requirement is particularly critical for SPIM techniques using submicron thick light sheets and high detection NA to achieve submicron axial resolutions because of the thin light sheet thickness and the narrow detection depth of the focus. For example, in Bessel SPIM or lattice light sheet microscopy, a half micron off focus of the excitation light from the detection focal plane is severe enough to decrease the spatial resolution and the signal to noise ratio (SNR) of the result significantly (Gao, 2015). Therefore, besides using an appropriate excitation light sheet in SPIM imaging, it is also important to keep the excitation light sheet in focus at all times to ensure the desired 3D imaging ability of SPIM.

Unfortunately, the requirement of keeping the excitation light sheet in focus is often violated in the practice of SPIM imaging. The focus drift of the detection objective, usually caused by the temperature fluctuation of the imaging system, is a major problem that leads to off focus of the excitation light sheet in SPIM. In addition, SPIM is perhaps more sensitive to the detection focus drift for two more reasons. First, the focus drift of the detection objective couldn’t be self-compensated as that in microscopes based on the epi illumination configuration because of the separated sample illumination and fluorescence detection. Second, both objectives are usually submerged in the imaging buffer, so that the temperature variation of the imaging system often projects to the focus drift of both objectives immediately. The problem is especially disturbing when the imaging buffer needs to be heated upon the requirement of the specimen to be imaged. Therefore, a solution is required to keep the excitation light sheet in focus in SPIM live imaging, in which an essential requirement is be able to locate the axial position of the excitation light sheet in situ, so that a correction can be applied accordingly when it is off focus.

A point light source, such as a fluorescent particle, is usually required to determine the focal plane or the axial position of the excitation light sheet accurately, by which the light sheet axial position can be obtained by measuring either the intensity profile of the light sheet, the point spread function (PSF) of the microscope, or the fluorescence intensity of the point light source. However, a point light source is barely available in most biological specimens. Although similar methods may still be applied in SPIM using sharp features of the sample structure, such as punctate and filament structures, the results are inaccurate due to the coupling of the unknown sample structure. Furthermore, appropriate sharp features are still rare in most biological specimens and it is difficult to embed a sample feature selection process into an automatic process. For the reasons above, most methods developed to correct the focus drift in live microscopy, not specifically for SPIM, are based on the frequency content analysis of the acquired images. The common basis of these methods is that the collected images contain higher contrast and more fine details of the sample when the microscope is in focus (Oliva et al., 1999; Sun et al., 2004; Liu et al. 2007; Yazdanfar et al., 2008; Royer et al., 2016). However, these methods are also not accurate enough due to the coupling of the unknown sample structure into the analysis process.

To solve the problem, we developed a novel method to find the axial position of the excitation light sheet in SPIM on general biological specimens, by which the influence of the sample structure is suppressed. In the method, a patterned light sheet is generated to illuminate the specimen at a fixed plane and a widefield 3D image stack of the fluorescence emission pattern is collected. As the high frequency signal attenuates much faster than the low frequency signal when the patterned light sheet deviates from the detection focal plane, the axial position of the excitation light sheet can be determined by identifying the image plane at which the strongest signal intensity at the modulation frequency is observed. The off focus of the excitation light sheet is corrected thereafter by resetting the offset position of either the excitation light sheet or the detection objective. In this article, we investigate the mechanism of the method and evaluate its performance using a Lattice light sheet microscope (Chen et al., 2014). We show the method gives the axial position of the excitation light sheet with hundreds of nanometer accuracy in seconds.

## Theory

The operation procedure of the method is shown in Fig. 1. A patterned excitation light sheet that contains a specific modulation frequency is parked at a fixed sample plane, while the detection objective is scanned for a few micron distance at a fixed step size to take a widefield 3D image stack of the fluorescence emission pattern. Next, 2D Fourier transform is applied to the acquired 3D image stack plane by plane. The ratio between the modulated DC component produced by the patterned light sheet and the original DC component at each image plane is calculated to determine the axial position of the excitation light sheet. The excitation light sheet is in focus at the image plane where the maximal ratio is observed, and the influence of the sample structure is nearly eliminated due to the dividing operation. Finally, the off focus of the excitation light sheet is corrected by resetting the offset position of either the excitation light sheet or the detection objective. The same operation can be repeated periodically to keep the excitation light sheet in focus for a long period of time.

**Figure 1.**
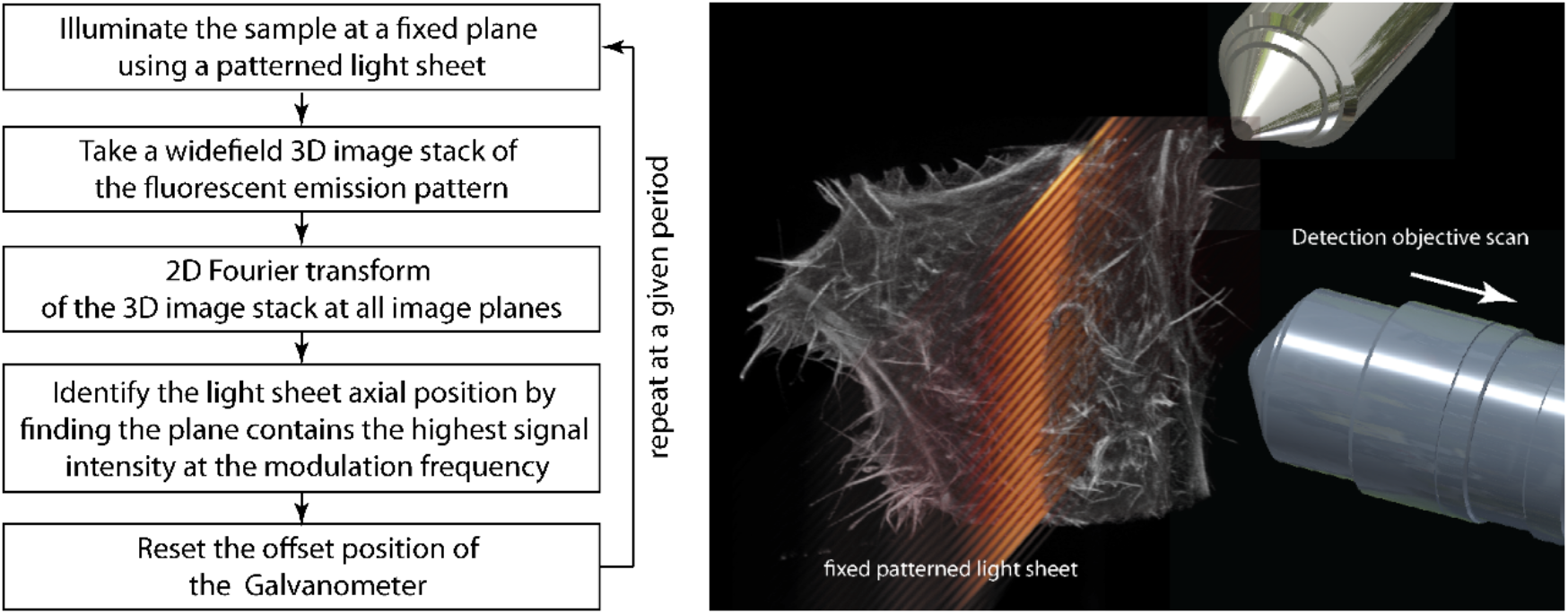
The operation procedure of the presented method. A patterned light sheet is parked at a fixed sample plane, while a 3D widefield image stack of the fluorescence emission pattern is collected and used to find the axial position of the excitation light sheet.

The concept of the method can be understood as follows. The patterned light sheet creates a fluorescence emission pattern that contains a specific spatial frequency in the modulation direction, in which the relative intensity of the modulated signal at the modulation frequency to the non-modulated signal in mainly determined by the illumination pattern. During the acquiring process of the 3D widefield image stack of the fluorescence emission pattern, the emission pattern is focused and defocused as the detection focal plane scanned across the excited sample plane. Because the high frequency signal attenuates faster than the low frequency signal when the emission pattern becomes off focus in widefield detection (Neil et al., 1997), the ratio between the modulated signal and the non-modulated signal is the maximum when the light sheet is in focus.

The following paragraph explains the mechanism of the method more accurately. The collected 3D image stack is the convolution of the microscope PSF and the produced fluorescence emission pattern. The relationship can be expressed as *D*(*r*) = *H*(*r*) ⊗ *E*(*r*), in which *D*(*r*) is the recorded image, *H*(*r*) is the PSF of the microscope, *E*(*r*) is the fluorescence emission pattern, and *r* denotes the spatial coordinates (*x, y, z*) in real space. Meanwhile, the fluorescence emission pattern can be described as *E*(*r*) = *S*(*r*)*I*(*r*), where *S*(*r*) represents the fluorescent labeled specimen, and *I*(*r*) is the intensity profile of the patterned light sheet parked on the specimen.

In SPIM, the intensity profile of a patterned light sheet can be described as the sum of multiple modulation harmonics:

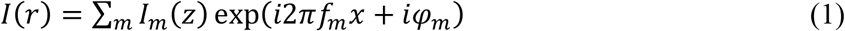

where *I_m_*(*z*), *f_m_* and *φ_m_* are the intensity, frequency and phase of each harmonic respectively (Gao et al., 2014), which are determined by how the patterned light sheet is generated. By substituting *I*(*r*) with Eq. 1, it can be obtained that

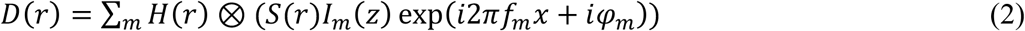

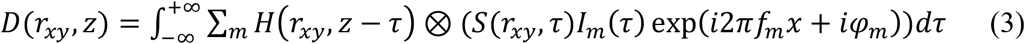

Therefore, the 2D Fourier transform of each image plane takes the form

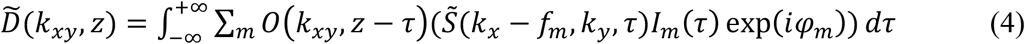

In which *O*(*k_xy_, z*) is the optical transfer function of *H*(*r_xy_, z*), and *k_xy_* defines the lateral coordinate in frequency space. As a result, the ratio between the modulated DC component by the *m*^th^ harmonic and non-modulated DC component at each image plane is given by

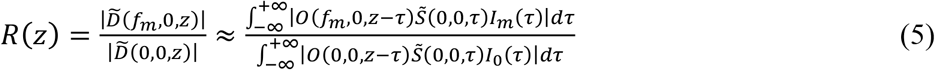

As long as the sample structure is roughly uniform, the equation can be approximately simplified as

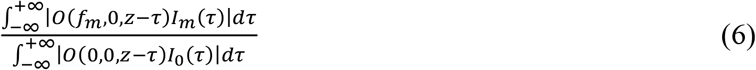

Obviously, the ratio *R*(*z*) reaches the maximum when the patterned light sheet is in focus, because *O*(0,0, *z*) is a constant, and both *O*(*f_m_*, 0, *z*) and *I_m_*(*z*) are the maximal at *z* = 0.

## Results

We evaluated the performance of the method by imaging fixed HeLa cells labeled with Vybrant CM-Dil on a lattice light sheet microscope. In lattice light sheet microscopy, a coherent Bessel beam array, which contains several modulation harmonics, is either dithered to illuminate the sample in the sheet scan mode, or scanned in discrete steps to illuminate the sample in the structured illumination mode.

At first, a coherent Bessel beam array generated with a 0.55 NA_od_, 0.5 NA_id_ annulus was parked at an arbitrary plane of the selected cell. The excitation objective was scanned for a 20 μm distance at a 150 nm step size to collect a 3D image stack. Fig. 2(a) and 2(b) show the selected image planes where the patterned light sheet is in focus and one micron off focus. Next, 2D Fourier transform was applied to each image plane. Fig. 2(c) and 2(d) show the corresponding 2D Fourier transform of the selected image plane in Fig. 2(a) and 2(b). It can be observed clearly that the modulation signal is much stronger when the light sheet is in focus in both real space and frequency space. The ratio *R*(*z*) was calculated and plotted against the axial position of the image plane as shown in Fig. 2(e) and 2(f). As expected, the maximal modulation intensity was observed when the excitation light sheet is in focus.

**Figure 2.**
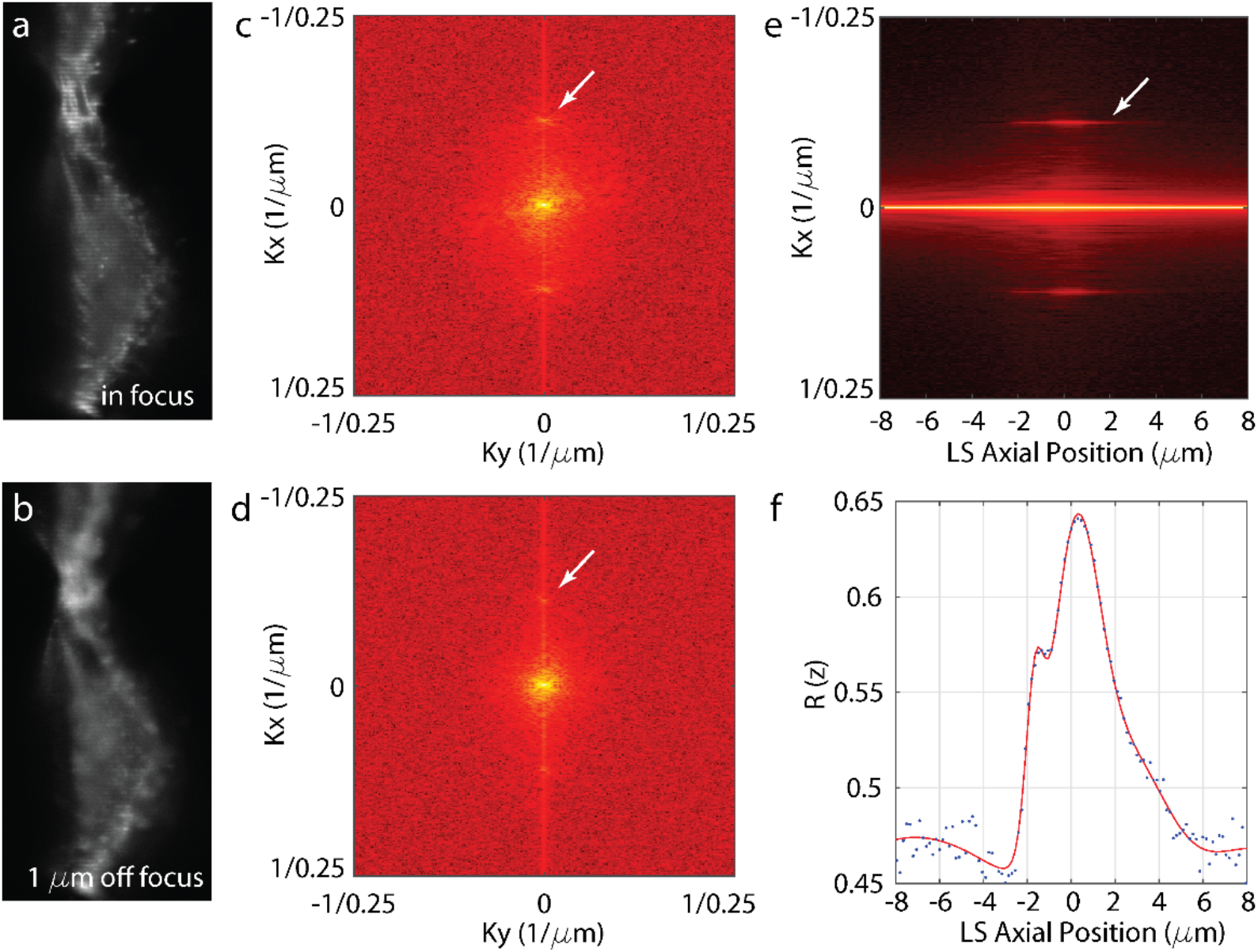
The axial position of the excitation light sheet is determined by identifying the image plane contains the strongest signal intensity at the modulation frequency. (a) The image plane at which the patterned light sheet is in focus. (b) The image plane at which the patterned light sheet is one micron away from the detection focal plane. (c) The 2D Fourier transform of the image plane in a. (d) The 2D Fourier transform of the image plane in b. (e) The intensity of the crossline at *k_y_* = 0 of the 2D Fourier transform of all image planes. (f) The relative intensity of the modulated DC component to the non-modulated DC component at different image planes. The ratio reaches the maximum when the light sheet is in focus. (A gamma value of 0.1 is applied in c-e to enhance the visibility of the modulation signal).

Next, we evaluated the long term focusing stability of the microscope using the same cell by repeating the same procedure described above every 10 minutes for ~12 hours. The results show that the light sheet drifted about ~2.5 μm according to the detection focal plane during the period (Fig. 3(a)), which is likely due to the temperature variation of the environment, despite the heating system of the microscope was running continuously. Fig. 3(b) and 3(c) show the relative modulation intensity *R*(*z*) of the collected 3D image stack at the first hour and the tenth hour, by which the axial position of the excitation light sheet can be located and the drifting can be observed clearly. Fig. 3(d) and 3(e) show that the imaging quality degraded quickly if the off focus of the excitation light sheet is not corrected.

**Figure 3.**
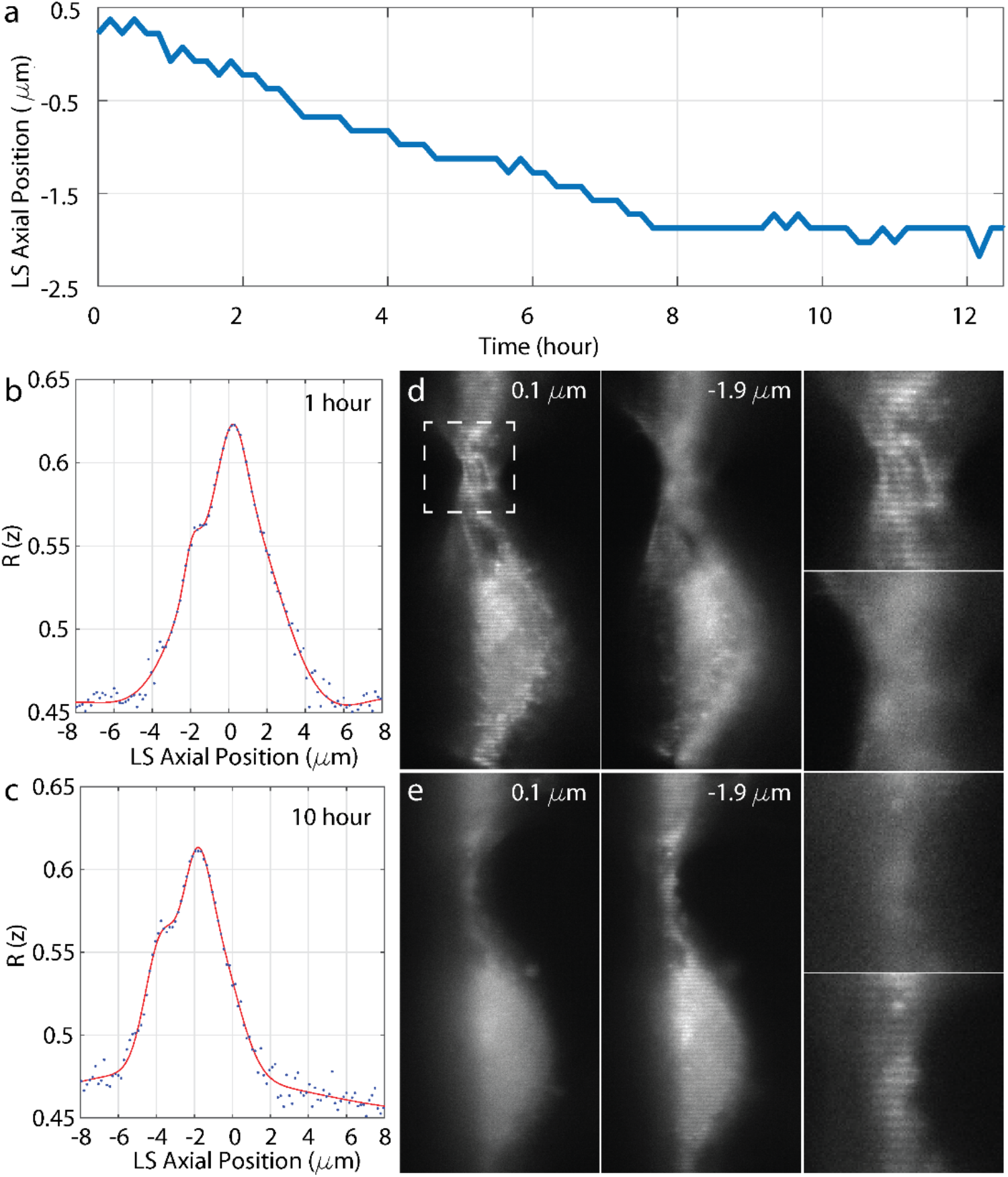
Monitoring the focusing stability of a lattice light sheet microscope using the presented method. (a) The drift of the excitation light sheet according to the detection focal plane in ~12 hours. (b-c) Locate the axial position of the excitation light sheet at the first hour and the tenth hour time points using the presented method. (e-f) The comparison of the same two image planes at the above two time points where the light sheet is in focus and off focus due to the uncorrected focus drift.

## Discussion

In summary, we developed a method to find the axial position of the excitation light sheet in SPIM on general biological specimens so that a correction can be applied based on the result to keep the light sheet in focus. We discussed the mechanism of the method and confirmed its ability on a lattice light sheet microscope by imaging HeLa cells labeling cell membrane. The method has several major advantages. First, the method is software based, and it can be implemented in an automatic process easily. Second, it can be applied to biological specimens without requiring special features, so that the method is applicable on most biological specimens. Third, it only requires dozens of 2D images to locate the axial position of the excitation light sheet, which is photon efficient in general, as the excitation light is highly confine in SPIM. It not only returns the result accurately in seconds, but also avoids the waste of the limited photon budget.

Besides the lattice light sheet microscopy technique used to demonstrate the method, the method can be implemented in any other SPIM technique that is capable to create a patterned light sheet, such as the SPIM techniques that create the excitation light sheet using a scanning beam or beam array (Keller et al., 2008, 2010; Planchon et al., 2011; Gao et al., 2012). Additionally, although the modulation frequency of the patterned light sheet is not critical, the method works the best when the modulation frequency is the half of that of the detection lateral diffraction limit, at which the widefield detection has the best axial resolving ability, so that the variation of the modulation signal intensity is the most sensitive to the defocusing of the patterned light sheet. Finally, the described method assumes that the labeled sample structure is roughly uniform. Therefore, it could fail when the sample is only sparsely labeled, e.g. the sample is a fluorescent particle. Fortunately, such situation is rare in most biological specimens, and other methods developed previously can be used instead to find axial position of the excitation light sheet.

## Declaration of competing Interests

A US provisional patent regarding the described technique was filed on behalf of L.G. by 3I Inc. on 05/04/2017.

